# vRhyme enables binning of viral genomes from metagenomes

**DOI:** 10.1101/2021.12.16.473018

**Authors:** Kristopher Kieft, Alyssa Adams, Rauf Salamzade, Lindsay Kalan, Karthik Anantharaman

**Affiliations:** Department of Bacteriology, University of Wisconsin–Madison, Madison, WI, USA; Microbiology Doctoral Training Program, University of Wisconsin–Madison, Madison, WI, USA; Computation and Informatics in Biology and Medicine, University of Wisconsin–Madison, Madison, WI, USA; Department of Medical Microbiology and Immunology, University of Wisconsin–Madison, Madison, WI, USA; Department of Medicine, University of Wisconsin–Madison, Madison, WI, USA

## Abstract

Genome binning has been essential for characterization of bacteria, archaea, and even eukaryotes from metagenomes. Yet, no approach exists for viruses. We developed vRhyme, a fast and precise software for construction of viral metagenome-assembled genomes (vMAGs). vRhyme utilizes single- or multi-sample coverage effect size comparisons between scaffolds and employs supervised machine learning to identity nucleotide feature similarities, which are compiled into iterations of weighted networks and refined bins. Using simulated viromes, we displayed superior performance of vRhyme compared to available binning tools in constructing more complete and uncontaminated vMAGs. When applied to 10,601 viral scaffolds from human skin, vRhyme advanced our understanding of resident viruses, highlighted by identification of a Herelleviridae vMAG comprised of 22 scaffolds, and another vMAG encoding a nitrate reductase metabolic gene, representing near-complete genomes post-binning. vRhyme will enable a convention of binning uncultivated viral genomes and has the potential to transform metagenome-based viral ecology.

## Main

Viruses and bacteriophages (collectively termed viruses) are pervasive members of essentially all ecosystems. Viruses form a continuum of symbiotic interactions with their hosts, from lethal parasitism to essential mutualism^1–3^. These interactions are known to impact biogeochemical and nutrient cycling processes, human health, infrastructure and industries, and ecosystem community dynamics^4–7^. As a result of the rising interest in viromics, the previously unknown members of the virosphere, the range in the encoded genetic potential of viruses, known viral diversity, and limits of viral genome sizes have been continuously expanding^8–12^.

Metagenomic sequencing can be a mechanism to identify, recognize, understand, and even harness the information encoded on viral genomes. Most metagenomes will assemble into many short fragments (scaffolds or contigs) representing partial genome sequences. The process of binning is employed to group scaffolds into a putative genome, termed a metagenome-assembled genome (MAG). With the information encoded by a MAG, rather than individual scaffolds, stronger inferences of metabolic potential, phylogenies, taxonomy, and community interactions can be generated^13^.

Many software tools have been developed for binning bacterial, archaeal, and eukaryotic metagenomic scaffolds into MAGs^14–23^. These tools employ a wide range of methodologies, mainly focusing on tetranucleotide frequencies and read coverage variance comparisons between scaffolds. A significant portion of these tools, most of which are tailored to bacteria and archaea, also rely on identifying microbial single copy genes to inform the construction of bins along with completeness and contamination estimates.

Conversely, viral scaffolds are typically not binned. Handling complex and often enigmatic viral scaffolds in metagenomes often poses computational challenges unique from microbes. One justification to not bin viruses is that their genomes are small relative to cellular organisms and the assumption that most scaffolds represent the majority, or the entirety, of an identifiable genome. For dsDNA viruses, the target of most viral metagenomes, genome sizes will have a general range of 20 kb – 200 kb, with the largest of viruses being 500 kb – 2000 kb. Since the majority of scaffolds in most assembled metagenomes are below 20 kb in length, it can be estimated that a single scaffold likely will not represent an entire viral genome. In fact, benchmarks have shown that viruses often do not assemble into a single scaffold^24,25^. Another difficulty with binning viral genomes is that viruses do not encode universal single copy or marker genes, making a standardized approach for all viruses difficult to create.

Despite the abundance of tools for binning bacteria and archaea, no tool currently exists for binning viruses. Here, we present vRhyme, a software tool that incorporates supervised machine learning based classification of sequence feature composition as well as read coverage effect size comparisons to optimize the binning of viral genomes from metagenomes (viral MAGs, or vMAGs). vRhyme leverages unique features of viral genomes, including overcoming the lack of single copy genes by scoring protein redundancy based on the observation that viruses seldom encode redundant genes. vRhyme is capable of binning viruses from diverse families, host and source environment affiliations, varying states of genome fragmentation, and wide ranges of genome lengths. In benchmarking vRhyme, we show that it is fast, inclusive, and accurate in binning viral scaffolds, with low computational demands, in synthetic and natural metagenomes compared to other binning software. When applied to human skin metagenomes, we show that vRhyme enabled a more comprehensive analysis of shared viruses and viral features across a cohort of individuals, and likely better recapitulated natural systems. vRhyme is implemented in Python and is freely available for download at https://github.com/AnantharamanLab/vRhyme.

## Results

### vRhyme overview and workflow

The vRhyme workflow is done in five steps: read coverage processing, sequence feature extraction, supervised machine learning, iterative network clustering, and bin scoring (**Figure 1**). The base input to vRhyme are the assembled scaffolds or contigs to be binned (hereafter scaffolds) with a set minimum size of 2 kb. For optimal results, only virome scaffolds or predicted virus scaffolds should be used as input, though vRhyme can function with the input of an entire metagenome. An initial dereplication step to remove redundant input scaffolds is optional. Next, scaffolds are compared pairwise by read coverage composition per sample, which is a proxy for relative abundance. vRhyme performs optimally with an input of multiple samples (i.e., coverage files) for more robust coverage co-occurrence estimations, but it will function with a single sample input with a minor decrease in performance. Statistically dissimilar scaffolds by coverage composition are screened out and the remaining potential pairs are compared by nucleotide feature similarity. Seven total nucleotide and gene features are used to classify pairs as similar versus dissimilar using two supervised machine learning models (decision trees and neural network). Following this step, potential connections are made between scaffolds based on similarity in read coverage and nucleotide features. These connections are used to create weighted networks that are further refined into genome bins using KMeans clustering. The entire process of read coverage comparison, nucleotide feature machine learning and weighted network refinement is performed over several *binning iterations* in parallel. vRhyme has 15 built-in presets of thresholds for Cohen’s *d*, machine learning model probabilities, and network edge weights. The number of presets used is equivalent to the number of binning iterations completed. A list of all presets and their hierarchy can be found in **Supplementary Table 1**. Each bin within all binning iterations is scored according to protein redundancy, a proxy for contamination, and the best binning iteration by sequences binned, bins generated, and redundancy metrics is selected. The bins within this best binning iteration are reported along with relevant metadata, including number of members and total protein redundancy. Alternative binning iterations are likewise saved if manual inspection and selection of a different iteration is desired.

**Figure 1.**
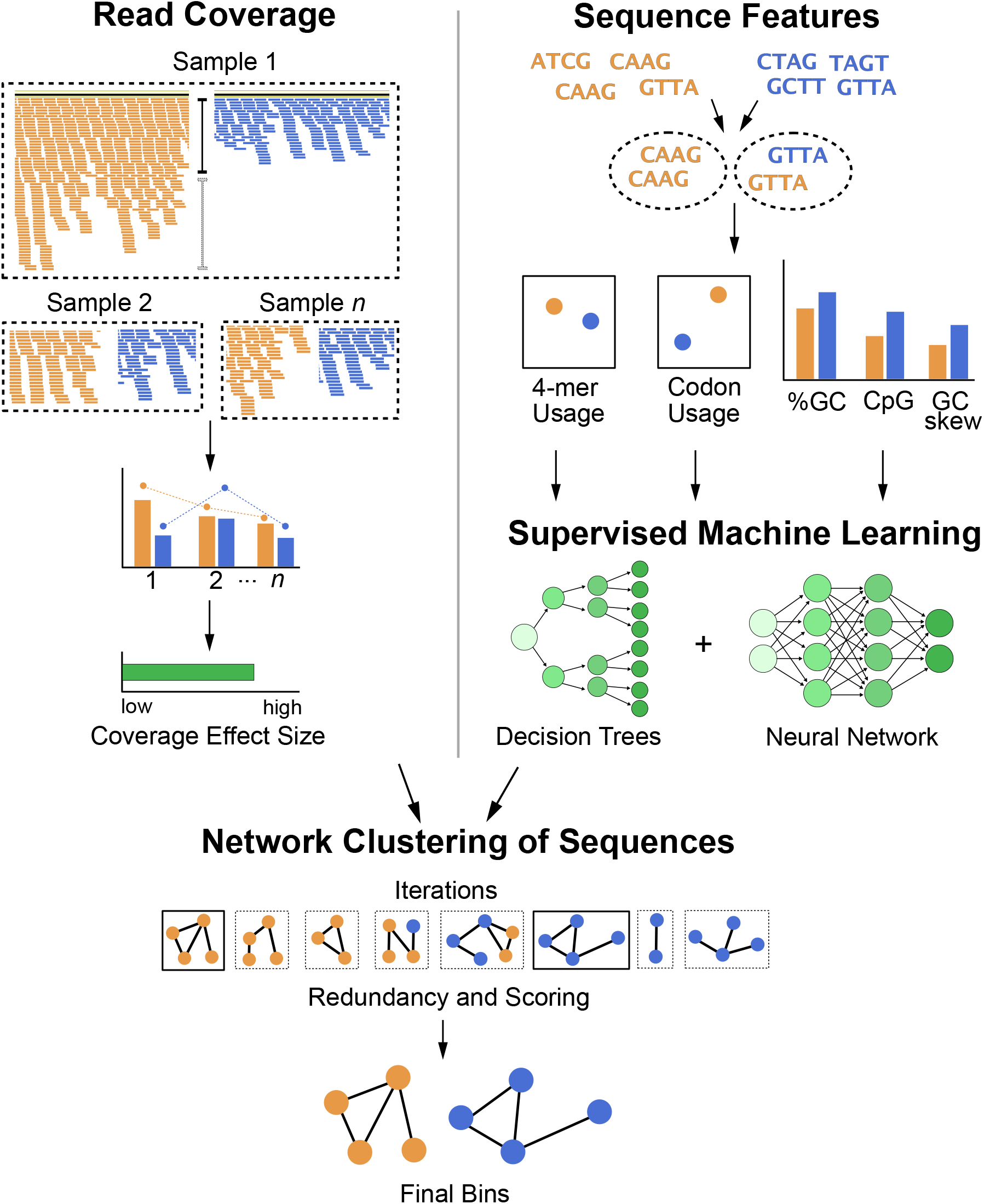
Flowchart of vRhyme workflow and methodology. Scaffolds are compared pairwise by read coverage effect size differences using single or multiple samples (top-left), followed by sequence feature distance comparisons (top-right). Multiple iterations of network clustering of putative bins are generated with edge weights representing normalized coverage effect size and supervised machine learning probabilities of sequence feature similarity (center). The bins are refined by KMeans clustering, and the best set of bins from a single iteration are identified after identifying protein redundancy and scoring (bottom).

### Assessment of binning quality

To evaluate vRhyme, we first benchmarked vRhyme against reference datasets and compared the performance to several available binning tools, all of which are built for microbes. Many binning tools and wrapper software were not suitable for viral binning due to reliance on microbial single copy genes. We were able to successfully compare vRhyme to MetaBat2^15^, VAMB^16^, CONCOCT^26^, and BinSanity^22^ on nine datasets curated from metagenomic data (see Methods). The nine datasets were comprised of 999 non-redundant and putatively complete viral genomes that were split 4,554 sequence fragments of varying lengths. Although these fragments were derived from datasets not represented in the machine learning training dataset, we first verified that the fragments were distinct and would not result in a bias associated with an overfitted machine learning model. Based on BLASTn similarity at 70% identity, only ∼5% of the 4,554 fragments were represented in the machine learning model training dataset, with all but four of the represented fragments being from the same human gut dataset.

A total of 17 different evaluation metrics were used, including five traditional metrics for recall, precision, accuracy, specificity, and F1 score (**Figure 2**). The five traditional metrics were calculated according to the true positive, true negative, false positive, and false negative rates of binning fragments together from the same or different source genomes (**Supplementary Table 2a**). Note that the machine learning models were not benchmarked individually since performance is measured based on the entire pipeline. vRhyme yielded the highest F1 score, the harmonic average of precision and recall, with an average of 0.87 across all nine datasets. MetaBat2 and VAMB performed equally with F1 scores of 0.81 and 0.82, respectively, but importantly VAMB only successfully binned three of the nine datasets due to input size requirements. vRhyme likewise yielded the highest, or equal to highest, average precision (0.94), accuracy (0.90), and specificity (0.96). Compared to MetaBat2 and VAMB, vRhyme likewise yielded the greatest average recall (0.80). CONCOCT and BinSanity yielded the greatest average recall values (0.96 and 0.91, respectively) but at the expense of precision (0.45 and 0.44, respectively). At least for viral genomes, CONCOCT and BinSanity were found to not be suitable binning options. VAMB had suitable performance on the three datasets with enough input sequences, but VAMB is likely not an option for many applications of binning viral genomes due to requiring many input sequences (e.g., tens of thousands^16^) for optimal performance. Based on these metrics, vRhyme performed exceptionally in binning viral genomes but did not considerably improve on the performance of MetaBat2.

**Figure 2.**
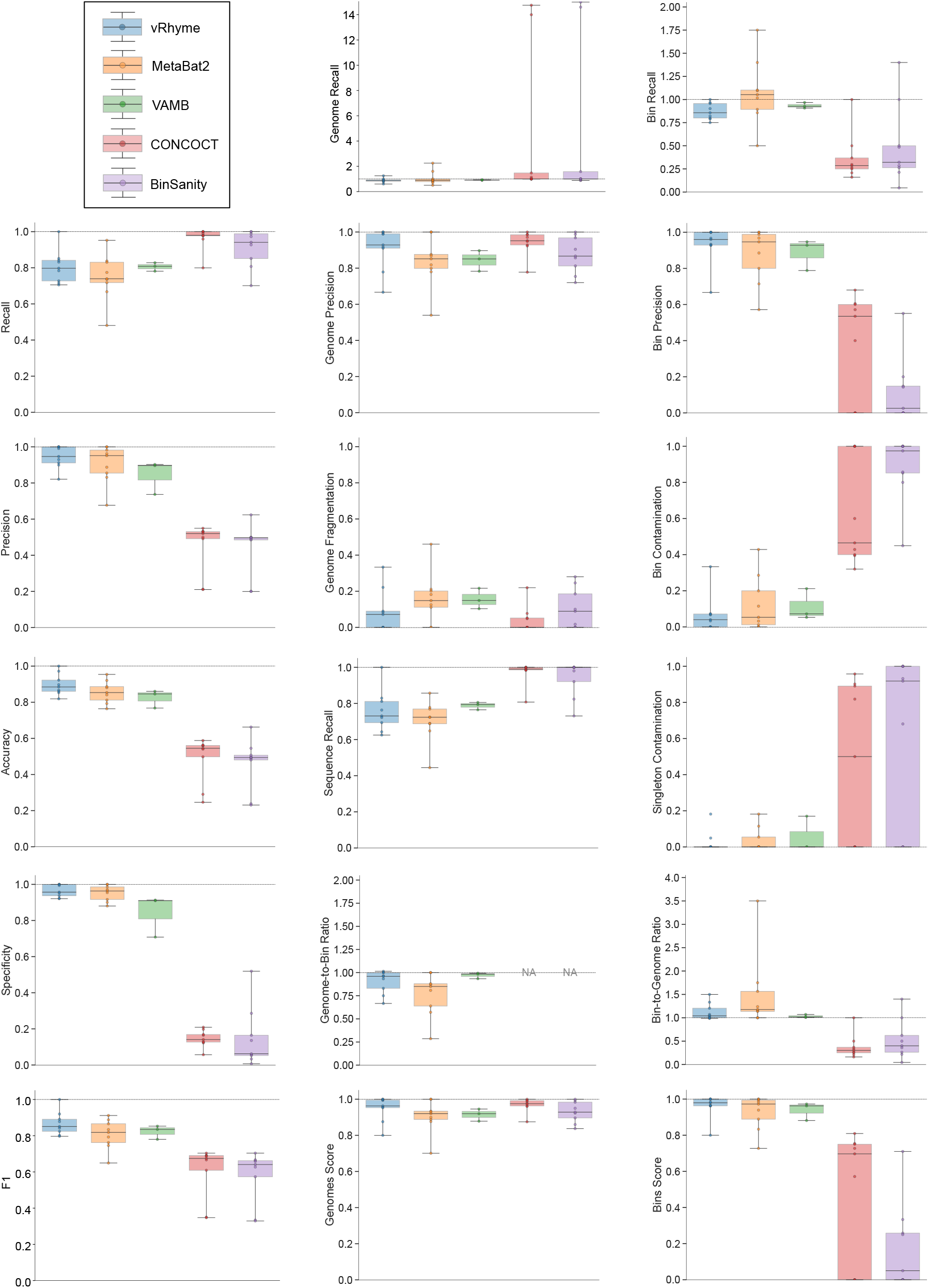
Benchmarking performance metrics of vRhyme compared to MetaBat2, VAMB, CONCOCT and BinSanity. Each boxplot represents the results of 9 different datasets, except for VAMB in which three datasets are shown. In total, 999 non-redundant genomes artificially split into 4,554 sequence fragments are shown. For some plots, a dotted line is shown at 1.0 to indicate optimal performance. CONCOCT and BinSanity are not shown on the Genome-to-Bin Ratio plot for better visualization; each yielded an average ratio greater than 2.0.

The remaining 12 evaluation metrics were calculated according to complete genomes and individual bins. These included evaluating if genomes were placed into a single or separate bins, and if bins contained fragments from a single or multiple source genomes. These metrics were better able to show the distinct performance of vRhyme compared to the other tools (**Supplementary Table 2b**). Namely, vRhyme was better able to reduce the following: placement of genomes into separate bins, placement of fragments from multiple source genomes into a single bin, and binning circular scaffolds representing entire genomes. Importantly, this was not at the cost of reduced fragment recall by vRhyme. To combine these metrics, we created a genome score and bin score that considered recall and precision as a substitution for F1 score. For genome scores and bin scores, respectively, vRhyme (0.89 and 0.96) outperformed, or was equivalent to, MetaBat2 (0.77 and 0.93) and VAMB (0.90 and 0.93). Again, it is important to note that VAMB only successfully binned three of the nine datasets. For CONCOCT and BinSanity, genome scores (0.74 and 0.70, respectively) and bin scores (0.48 and 0.18, respectively) reflected the propensity to “over bin” distinct genomes together into one bin.

Furthermore, we evaluated how well vRhyme bins compare to the input, unfragmented genomes. First, using CheckV^27^ we show a distinct change in genome completeness estimation in the binned versus unbinned sequence fragments. vRhyme was able to recapitulate the completeness of the input genomes (**Figure 3a**). This is supported by a similar observation in the length of the input genomes versus the bins (**Figure 3b**). Moreover, we estimated the taxonomy of the input genomes, fragments, and binned vMAGs. We identified a distinct decrease in the ability to identify taxonomy of the fragments, which were rescued by binning (**Figure 3c**). The identifiable difference in the vMAGs is a lack of Microviridae. Yet, this is to be expected since the small genome size of Microviridae (<10 kb) typically results in near-complete scaffolds that appropriately remain unbinned. Finally, we evaluated whether vRhyme could distinguish the source scaffolds. To do this, each of the nine datasets were binned, but the scaffolds were not fragmented. The expected result is that none of the circular scaffolds should bin together. Although vRhyme did bin 11% of the whole scaffolds, it was a marked improvement on MetaBat2 which binned 65% (**Figure 3d**).

**Figure 3.**
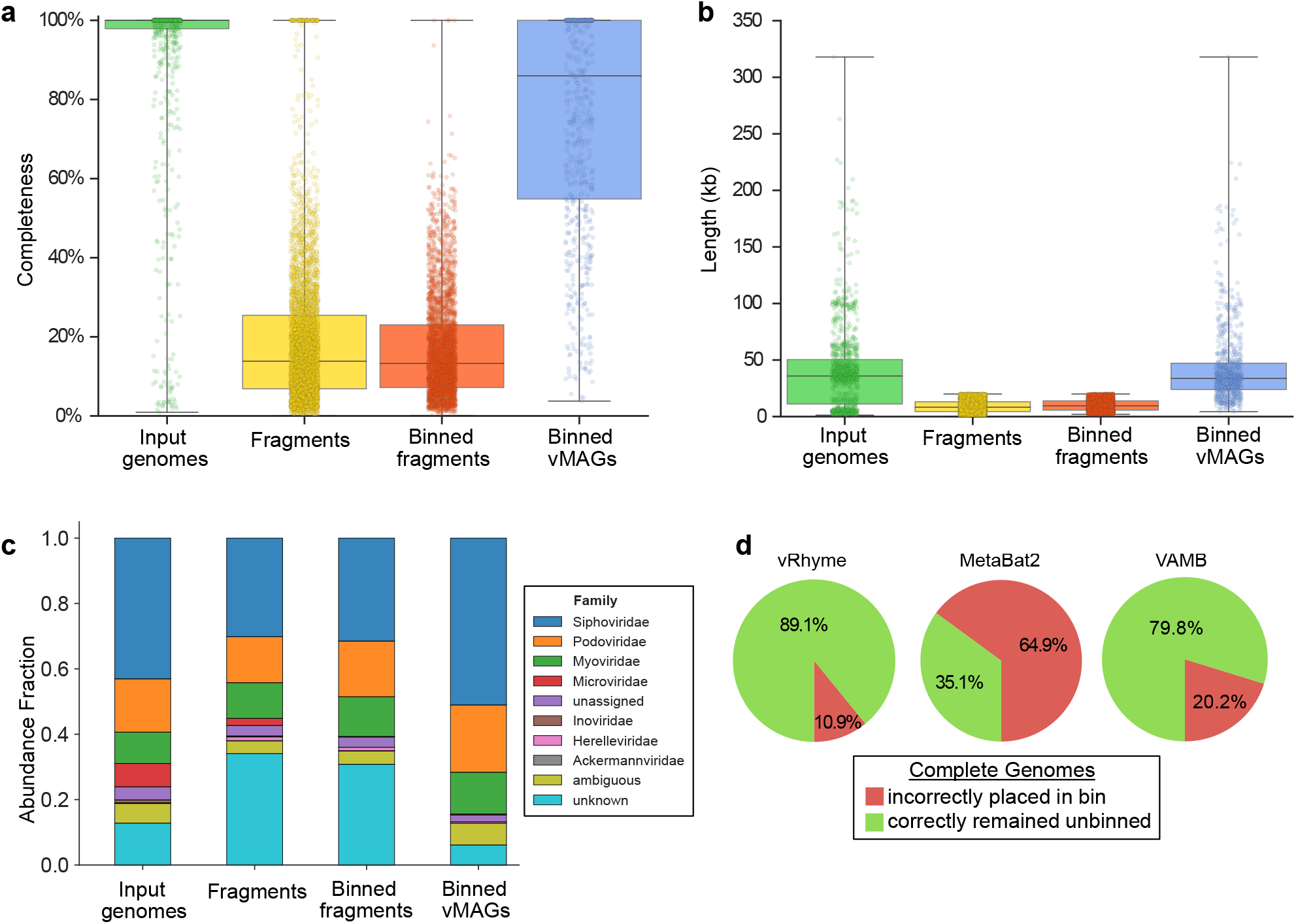
Impact of binning with vRhyme on the benchmarking datasets. For **a-c**, the putatively complete unsplit input genomes, generated sequence fragments, binning sequence fragments, and vRhyme bins (vMAGs) are compared. **a**, Estimation of genome completeness using CheckV. **b**, sequence or vMAG nucleotide length. For **a-b**, each dot represents a single sequence or vMAG. **c**, Estimation of taxonomy at the family level using a custom analysis script. “unassigned” represents a taxonomic classification to a group with an unassigned family, “ambiguous” represents equal assignment to multiple families (typically Caudovirales), and “unknown” represents the inability to make a prediction. **d**, Evaluation of vRhyme, MetaBat2 and VAMB for the binning of complete genomes. The expectation is that complete genomes should remain unbinned as vOTUs or UViGs.

### Discovery of vMAGs in human skin metagenomes

To demonstrate the ability of vRhyme to aid metagenome analyses and discovery, we applied vRhyme to 270 human skin metagenomes^28^. Viruses were predicted from a cohort of 34 individuals with eight body sites (*Af, Al, Ba, Na, Oc, Tw, Um*, and *Vf*) sampled per individual (see Methods). From all individuals, 10,601 viral scaffolds were identified and binned, across eight different body sites individually, into a total of 849 vMAGs representing 2,794 viral scaffolds. Although bins with redundant proteins may in fact be a single genome, we ignored all vMAGs with greater than one redundant protein for analysis to yield 762 vMAGs representing 2,413 viral scaffolds, leaving the remaining 8,188 as discrete viral scaffolds (**Supplementary Table 3**) (**Figure 4a**). The bins were comprised of an average of 3.2 scaffolds each. In total we identified seven bins, representing separate body sites, that were present across at least 30 individuals (**Figure 4b**). Two bins of unique characteristics were identified and examined in detail.

**Figure 4.**
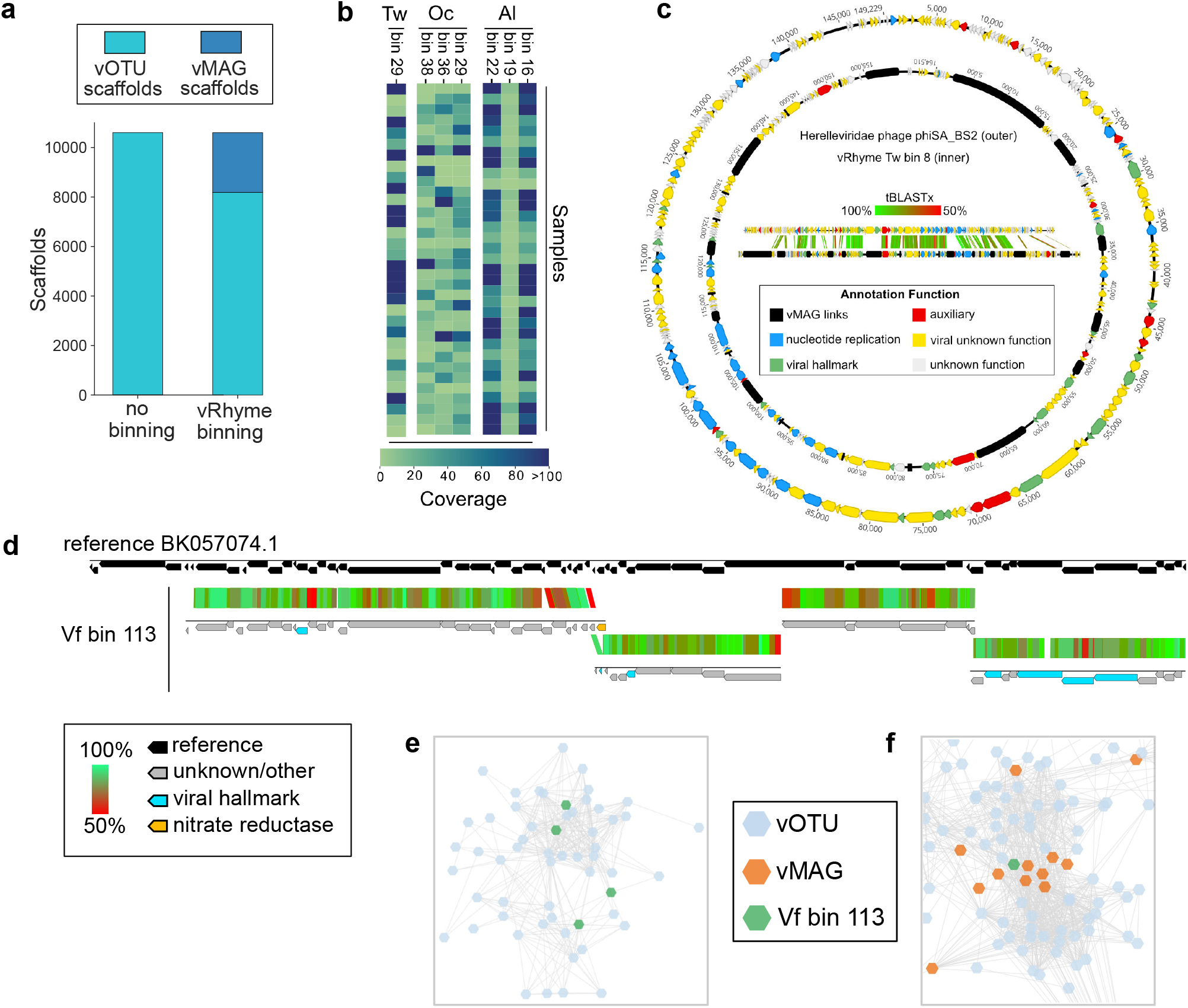
Binning improves and expands the analysis of viruses from human skin. **a**, Comparison of the number of original viral scaffolds identified across all individuals before and after binning. **b**, Heatmap of coverage for the seven common bins per individual. **c**, Genome visualization and alignment of Herelleviridae reference phiSA_BS2 (outer) and Tw bin 8 (inner). Each arrow represents a predicted open reading frame and black bars are artificial connections between vMAG scaffolds. **d**, Alignment of vRhyme Vf bin 113 to the closest reference virus Siphoviridae isolate ctiXA4 (BK057074.1). Each of the four scaffolds were independently aligned by tBLASTx similarity. The *narG* AMG is labeled in yellow and viral hallmark annotations are labeled in light blue. **e**, Representative cluster from all input viral scaffolds generated by vConTACT2, with the four Vf bin 113 scaffolds labeled in green. There are no connections between any of the four green scaffolds. Each dot represents a single scaffold. **f**, Partial network from all vRhyme binned and unbinned viral scaffolds generated by vConTACT2, with vMAG bins labeled in orange and Vf bin 113 in green. For **e**,**f** complete network diagrams can be found in **Supplementary Figures 1 and 2**.

The first such bin contained 22 members (Tw bin 8), more than what would be expected for a viral bin, and aligned to a reference Herelleviridae phage (Staphylococcus phage phiSA_BS2) (**Figure 4c**). Herelleviridae infecting abundant *Staphylococcus* on the skin are likely to be highly relevant to skin ecology and disease^29^. Before binning, each of the 22 members were identified by CheckV as low-quality genome fragments with individual completeness estimations ranging from 1.8% to 7.1%. The fragments averaged 5.2 kb in length and ranged from 2.6 kb to 10.0 kb. After binning, the final bin was 115 kb in length and identified as a high-quality genome with 100% completeness by CheckV. The reference phage genome is 143 kb, suggesting the true completeness of the bin is likely 80% to 100%. All CheckV results for the skin metagenomes can be found in **Supplementary Table 4**.

The second bin of interest contained 4 members (Vf bin 113), with one encoding a nitrate reductase (*narG*) auxiliary metabolic gene (AMG) (**Figure 4d**). The *narG* was positioned as the last gene on a scaffold, and conventional approaches for AMG validation would suggest discarding the AMG as likely bacterial contamination. However, binning aided in the validation of the AMG as likely to be correct. The first line of evidence was the lack of any integrase or lysogenic viral signatures on any of the four binned scaffolds, suggesting the AMG is not from bacterial contamination resulting from host integration. Second, alignment of all four scaffolds to the nearest reference genome (Siphoviridae isolate ctiXA4) displayed that the AMG was situated at the intersection of two scaffolds within the genome rather than at a genome end. CheckV identified each member as low-quality with completeness values of 11.6% to 28.0% for the respective 7.4 kb to 16.8 kb scaffolds. The bin was estimated to be of medium-quality with a completeness of 74.9%, or 92% based on the length of the closest reference genome. Moreover, one of the four scaffolds lacked characteristic viral annotations to aid with manual inspection or analyses such as phylogeny, yet binning with the other scaffolds containing viral hallmark and nucleotide replication annotations was able to validate the scaffold as viral and place it in better genomic context for analysis. Therefore, binning was able to not only generate a more complete sequence, but also validate the presence of an understudied and ecologically important AMG. Using vConTACT2^30^, we clustered the bin with the complete binning results (low-contamination bins plus unbinned scaffolds) (**Figure 4e**) in addition to clustering all of the individual, unprocessed viral scaffolds (**Figure 4f**). Binning of the individual scaffolds placed all four scaffolds of the bin into a single cluster distinct from other groups, yet as anticipated none of the scaffolds of the bin were connected. Clustering of the binning results yielded more connections between scaffolds and vMAGs and better placed the bin within evolutionary and community relationship contexts. Complete vConTACT2 networks can be found in **Supplementary Figures 1 and 2**.

## Discussion

Binning viral scaffolds into vMAGs is uncommon, with most or all remaining as discrete virus operational taxonomic units (vOTUs) or uncultivated virus genomes (UViGs)^31^. We believe adopting a more genome-centric approach for UViGs will enable innovative discoveries, such as the construction of large or highly heterogenous viral genomes that often assemble into dissimilar fragments. Here, we have presented vRhyme and demonstrate that the “one scaffold, one virus” convention can skew interpretations of a virosphere and the interactions of its viral community members. To address this, vRhyme enables the binning of viral genomes into vMAGs using a virus-centric approach, unique from existing binning software, in an easy to use and reproducible command line tool.

In addition to performance benchmarks on artificial and real metagenomes, we evaluated the robustness of vRhyme by binning artificially fragmented NCLDV, megaphage, large eukaryotic virus, crAssphage, active and inactive integrated prophage, and microbial genomes (**Supplementary Information**). vRhyme was largely capable of precisely binning these unique and complex viral datasets. However, notable exceptions were difficulties with separating multiple inactive (non-replicating) prophages from the same host genome as well as binning non-viral genomes, though the latter was an anticipated limitation. Moreover, we displayed that vRhyme is efficient and likely precise in binning large and complex datasets using Global Ocean Virome 2 (GOV2)^32^ and agricultural soil viromes^33^ (**Supplementary Information**). In total, we hope that with the availability of vRhyme as a reliable binning tool, vMAG construction will become a common practice and adopted into existing frameworks of studying viral ecology, host associations, community interactions, evolution, and biogeochemical cycling.

To further evaluate the computational capabilities of vRhyme or potential restraints, we assessed the effect of the coverage calculation methods, the number of input coverage samples and the effect of user-modifiable parameters on performance, as well as the runtime, memory usage and reproducibility of binning (**Supplementary Information**). We found that vRhyme performs optimally with multiple input samples for more robust coverage variance comparisons, though the optimal value depends on how the dataset or metagenome was constructed. For example, a metagenome assembled from a single, standalone sample may perform suitably. As for modifying parameters, vRhyme likely will yield optimal results with the default settings due to the built-in binning iterations. Furthermore, the runtime of vRhyme for average sized viral datasets was on the scale of seconds. The GOV2 dataset, the largest dataset evaluated, finished in 93 minutes with 2.3 GB of memory using 15 CPU threads. Lastly, the methods employed by vRhyme allow it to be fully reproducible. Overall, we found the necessary requirements to be relatively low and even possible on personal laptop systems.

There are several important considerations in the binning of vMAGs that are unique from microbial MAGs. First, any viral scaffold not contained within a bin (vMAG) should be considered as a vOTU or UViG. This aligns with the “one scaffold, one virus” convention which is likely true for many viral genomes, especially circular and complete genomes. In the skin datasets presented here, ∼23% of the viral scaffolds were binned into low contamination vMAGs and the remaining ∼77% should still be utilized in analyses as discrete scaffolds. Second, an entire metagenome can be used as input to vRhyme, or viral binning in general, with the caveat that contamination of bins with non-viral sequences may be higher with the added advantage that fewer viral scaffolds may be missed. For example, many phage genomes are arranged in cassettes such that structural, nucleotide replication, lysis and auxiliary genes form distinct regions. If these regions were to assemble into separate scaffolds, virus identification may only identify a portion of the scaffolds, such as missing an auxiliary region, whereas binning may place them all together into a single vMAG. Third, accurate read coverage profiles are crucial for accurate binning. This is true for all binning software that depend on differential coverage and is especially true for distinguishing bins of integrated prophages from a single host population. vMAGs representing prophages generated by vRhyme will likely represent the greatest fraction of redundant, contaminated bins.

## Methods

### Coverage processing

The input for read coverage information is variable: paired or unpaired short reads, SAM alignment file, BAM alignment file, or a pre-calculated coverage table. For short reads input, reads will be aligned to input scaffolds using either Bowtie2^34^ or BWA^35^; Bowtie2 is run with the parameters --no-unal --no-discordant, the latter being for paired reads only, and BWA is run with the mem algorithm. All reads should be quality filtered before being used as input. The resulting SAM alignment file, or an input SAM alignment file, will be converted into BAM format using Samtools^36^. BAM alignment files, either generated by the vRhyme pipeline or as user input, will then be processed. As such, any input combinations of short reads, SAM or BAM alignment files are compatible. BAM alignment files, if not already provided as input, are sorted and indexed using Samtools.

The Python package Pysam (https://github.com/pysam-developers/pysam) is then used to fetch aligned records within sorted and indexed BAM alignment files for processing and coverage calculations. First, aligned reads are filtered according to the percent identity alignment, as calculated by the sum of the number of gaps *g* and the number of mismatches *m* in the alignment divided by the length of the alignment *I*. The default is a 97% identity alignment.

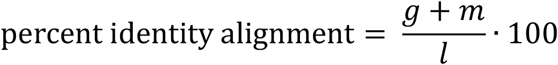

Aligned reads passing the set threshold are used to calculate the total coverage of each nucleotide base per scaffold, inclusive of bases with a coverage of zero. Finally, the coverage values at the terminal ends of scaffolds are masked to increase coverage fidelity by considering erroneous read alignment at partial scaffold ends. The default is to ignore all coverage values within the first and last 150 bp of the scaffold. The average and standard deviation of coverage per scaffold is calculated according to respective, individual base coverages. All alignment filtering and coverage calculations are handled natively within vRhyme. This final step yields a coverage table comprised of the average and standard deviation of coverage per scaffold per input sample. This coverage table, or a user-generated table of the same format, can be used as input for vRhyme in place of reads or SAM/BAM alignment files.

Next, scaffold coverages across all *k* samples are pairwise compared using the effect size of coverage differences. First, all average coverages are increased by a pseudo-count of 0.1 to avoid coverages of zero (pseudo-counts are excluded from coverage table). Effect size is calculated by the Cohen’s *d* effect size metric equation^37^. Cohen’s *d* is calculated as follows, where 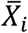 and 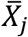 are average read coverages and σ_*i*_ and σ_*j*_ are standard deviations of the coverages for a scaffold pair *i* and *j*:

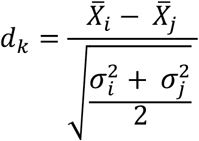

For each pairwise comparison, an effect size value *d*_*k*_ is generated per sample *k*. Values exceeding the effect size threshold, set by vRhyme presets, generate an additive penalty weight *p*. The average effect size across all samples 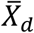, with any added penalties, is normalized to the number of input samples, yielding a normalized effect size *d′*, which considers higher statistical power to more sample comparisons:

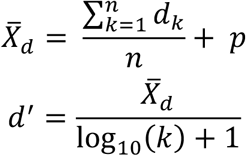

The normalized and penalized *d*^′^ values are compared to a normalized preset effect size threshold and all pairwise comparisons passing the set criteria are considered as co-occurring by coverage. Any scaffold not found to co-occur with another is discarded. For computational efficiency, a pre-filter is applied where only the best (i.e., lowest *d*^′^) *n* pairs per individual scaffold are retained, where *n* is ‘--max_edges’ multiplied by 3.

### Nucleotide processing

All co-occurring scaffolds by read coverage are compared by seven nucleotide content metrics. The pairwise distance calculations per metric are used as inputs to supervised machine learning models for classification. All nucleotide features and distances are calculated natively within vRhyme.

The first feature, codon usage (CU), is calculated from nucleotide open reading frames (i.e., genes). Predicted genes can be used as input, otherwise vRhyme will automate prediction using Prodigal^38^ (-m -p meta). In-frame trinucleotide counts *c* for each of the 64 codons *k* (step of 3 bases) along a scaffold are divided by the total count of observed codons. The final codon, if representing a stop, is ignored. Counts are inclusive of zero counts but exclusive of ambiguous (e.g., N) bases. The following yields a CU frequency vector *F*_*i*_ for each codon *k* in scaffold *i*.

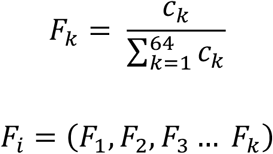

The next three features (GC content, CpG content, and GC-skew) are calculated per scaffold from individual scaffold bases. GC content *N*_.*gc*_ is calculated by the sum of all G and C bases, divided by the sum of all bases (A, T, C and G). CpG content *N*_*cpg*._ is calculated by the sum of all CG di-nucleotides per scaffold (step of 1 base) divided by the sum of all bases. GC-skew *N*_*skew*_is calculated by subtracting the total of C bases from the total G bases, divided by the sum of G and C bases.

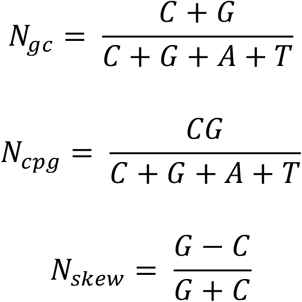

The last three features – relative tetranucleotide frequency (RTF), tetranucleotide usage deviation (TUD) and tetranucleotide zero’th order Markov method (ZOM) – are calculated from whole scaffold tetranucleotide frequencies (step of 1 base) of the forward and reverse strands^39^. A total of 136 possible tetranucleotides are considered after combining identical, reverse complement and palindromic sequences. Counts are inclusive of zero counts but exclusive of ambiguous (i.e., N) bases.

For RTF, all counts *t* for each of the 136 tetranucleotides *k* along a scaffold are divided by the total count of observed tetranucleotides. The following yields a tetranucleotide frequency vector *T*_*i*_ for each tetranucleotide *k* in scaffold *i*.

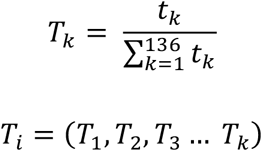

For TUD, expected nucleotide frequencies *E* are first calculated by dividing the count of each base *b* by the sum of all bases in the scaffold. Next, observed counts per base *O*_*b*_ per tetranucleotide *k* are calculated by the sum of each base inclusive of zero counts. For each unique tetranucleotide, expected frequencies per base are raised to the power of observed frequencies multiplied by two to yield a deviation value *D*_*b*_ per base. The deviation values for all four bases are multiplied the count of total observed tetranucleotides and the count of the given tetranucleotide to yield a TUD value per tetranucleotide. The following yields a TUD frequency vector *TUD*_*i*_ for each tetranucleotide *k* in scaffold *i*.

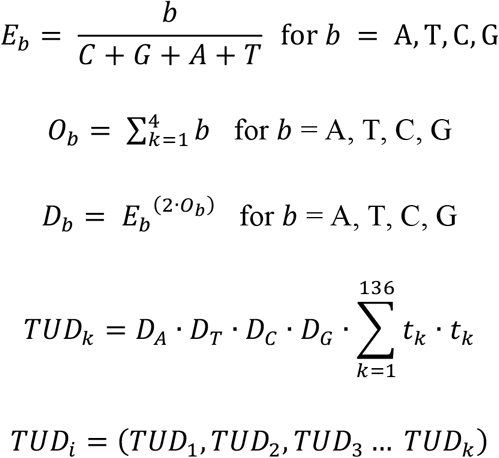

For ZOM, the same expected *E*_*b*_ nucleotide frequencies per base *b* are used. For each tetranucleotide *k*, the count *t* of the given tetranucleotide is divided by the product of each of the present tetranucleotide’s bases’ expected frequencies to yield a ZOM frequency vector *ZOM*_*i*_ for each tetranucleotide *k* in scaffold *i*.

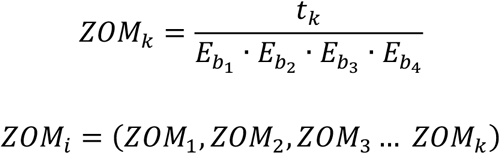

Pairwise distance calculations for GC, CpG and GC-skew are made by the absolute value difference in the respective metric’s content between two scaffolds. For example, the following is the pairwise distance *p*_*GC*_ in GC content between scaffolds *i* and *j*.

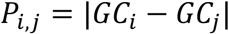

Pairwise distance calculations for CU, RTF, TUD and ZOM are made by cosine distances. For each value *v*_*i*_ and *v*_*j*_, corresponding to the same tetranucleotide *k*, in frequency vectors of scaffolds *i* and *j*, with vector averages of 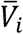 and 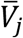, cosine similarity *S*_*i,j*_ is calculated. Cosine distances between two scaffolds are calculated for CU, RTF, TUD and ZOM individually.

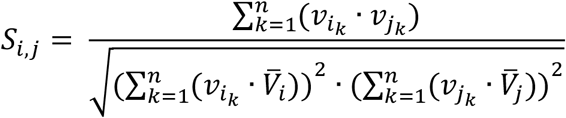

The result of distance calculations is a vector *M*_*i,j*_ of length seven for each pairwise comparison between scaffolds *i* and *j*.

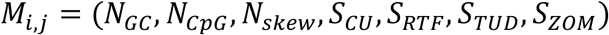

### Machine learning model training and testing

NCBI databases (RefSeq^40^ and Genbank^41^, release July 2019) were queried for “prokaryotic virus” and genomes greater than 10 kb in length were retained. In addition, the IMG/VR database (release July 2018)^42^ was downloaded, and sequences were limited to a length of 10 kb. For the IMG/VR dataset, VIBRANT^43^ (v1.2.1, -virome) and CheckV^27^ (v0.6.0) were used to obtain circular and/or complete sequences. The resulting NCBI and IMG/VR datasets were dereplicated by 95% identity using the method described here (--derep_only --derep_id 0.95 –frac 0.70 --method longest) and combined. The sequences representing complete genomes in the combined dataset were split into non-overlapping fragments of 15 kb with a minimum length of 10 kb. A total of 19,082 fragments were generated for training and testing machine learning models.

The machine learning models were generated based on the *M*_*i,j*_ vectors described above using the generated 19,082 genome fragments. Filtering of pairwise comparisons before training and testing was made according to vRhyme default parameters (--max_gc 0.20 --min_kmer 0.60). The pairwise comparison matrix was split 75:25 for training and testing, respectively. Fragment pairs were labeled as “same” or “different” for supervised machine learning according to if the paired fragments originated from the same or different source genomes. An equal number (69,632) of “same” and “different” pairs were used for training by randomly dropping excess “different” comparisons. For testing, a set of 38,685 “different” and 7,736 “same” pairs were used.

Scikit-Learn (v0.24.2)^44^ was used to generate machine learning models using a grid search approach to optimize parameters. Several models and algorithms were considered, including MLPClassifier, ExtraTrees, KNeighbors, SVC, Gradient Boost, Decision Tree and Random Forest classifiers. Iterative training and testing yielded MLPClassifier (alpha=0.001, beta_1=0.7, beta_2=0.8, hidden_layer_sizes=(5,25,50,75,100,100,75,50,25,5), learning_rate_init=0.0001, max_iter=1250, n_iter_no_change=15, tol=1e-08) and ExtraTreesClassifier (max_depth=10, max_features=7, n_estimators=1500) as the most robust.

### Machine learning and network processing

Each scaffold pair is classified by the two machine learning models separately to yield two probability values of “same”, one per model. The probability values are averaged to yield 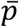. Any pair with 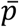 below the preset threshold is discarded. Then, *d*^′^ calculated previously for the pair is divided by 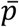 to yield a network edge weight *w*.

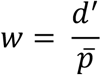

Any pair with *w* below the preset threshold is retained for network clustering. As before, for computational efficiency, only the best (i.e., lowest *c*) *n* pairs per individual scaffold are retained, where *n* is ‘--max_edges’. Weighted networks, representing unrefined bins, are created where each node is a scaffold and each edge is a weighted connection between paired scaffolds. Networks are refined using MiniBatchKMeans implemented in Scikit-Learn with the following parameters: n_clusters=*s*+1, batch_size=*h*, max_iter=100, max_no_improvement=5, n_init=5. Batch size *h* is 25% of the number of nodes with a minimum of 2 and maximum of 100. The number of clusters *d* is defined by the number of nodes with a clustering coefficient value below the preset constant 0.36 but not 0. For each node *i*, the clustering coefficient *U*_*i*_ is calculated as follows, where *L*_*i*_ is the degree of the node and *R*_*i*_ is the number of edges between the neighbors of *i*:

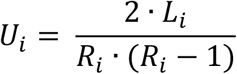

Refined networks are split into distinct, separate networks according to *s*. Here, each connected network represents a putative bin.

### Score processing

Each binning iteration is given a score *I* according to protein redundancy, total bins, and the number of scaffolds binned. To calculate protein redundancy, all proteins within a bin are clustered using Mmseqs2^45^ (linclust --min-seq-id 0.5 -c 0.8 -e 0.01 --min-aln-len 50 --cluster-mode 0 --seq-id-mode 0 --alignment-mode 3 --cov-mode 5 --kmer-per-seq 75). Any proteins clustered within a bin, excluding those along the same scaffold, are considered redundant. The iteration with the maximum score is selected as the final representative. *I* is calculated as follows:

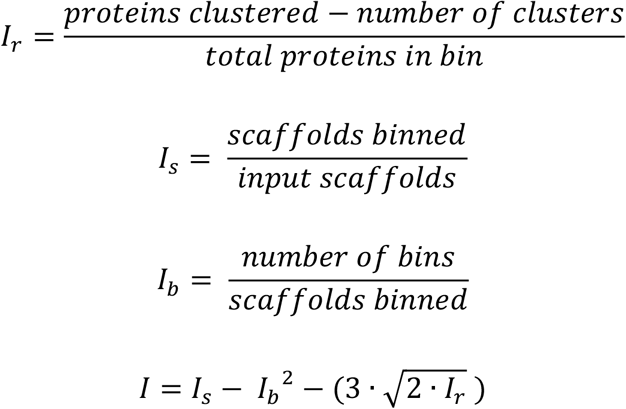

### Dereplication

vRhyme implements Nucmer^46^ and MASH^47^ for the dereplication of scaffolds. First, scaffolds are roughly grouped using MASH (sketch -k 31 -s 1000; dist) to reduce the pairwise comparison space. Next, all possible pairs of scaffolds within each resulting group are aligned using Nucmer (-c 1000 -b 1000 -g 1000). The longest scaffold in a pair with 100% identity over 100% coverage is taken as the representative. For all percent coverage calculations in dereplication, coverage is of the shortest scaffold. For ‘--method longest’ the longest scaffold in pairs meeting the percent identity and percent coverage thresholds is taken as the representative. For ‘--method composite’, scaffold pairs meeting the percent identity and percent coverage thresholds are joined over the region of sequence overlap to yield artificially chimeric scaffolds. Any alignments exceeding the sensitivity values for merging over complex alignments, such as low identity scaffold ends without overlap, are not joined. After scaffold pairs are joined, identical cycles of MASH, Nucmer and composite joining are completed until no further alignments are detected. For all methods, reverse complement sequence alignments are considered and adjusted accordingly.

### Performance validation datasets and metrics

Scaffolds used to benchmark performance were acquired from nine separate publicly available datasets spanning eight unique metagenomes from marine^48,49^, freshwater^50–52^, human gut^53^, and soil environments^54,55^. Details on the studies, scaffolds, reads, and accession numbers can be found in **Supplemental Table 11** Each dataset was processed separately. First, VIBRANT (v1.2.1) was used to predict viruses. From these viruses, VIBRANT and CheckV were used to identify circular scaffolds representing complete genomes. Next, scaffolds were dereplicated by 97% identity using the method described here (--derep_only --derep_id 0.75 --frac 0.70 --method longest). The non-redundant scaffolds were randomly fragmented into sequences ranging from 2 kb to 20 kb in length. A total of 999 scaffolds (i.e., putatively complete genomes) were used to generate 4,554 fragments. Full benchmarking was performed on the 4,554 fragments and validation of complete genome binning was performed on the 999 scaffolds representing complete genomes.

Since the circular scaffolds (sources) were estimated to be complete genomes, any of the fragments originating from the same source were expected to create a single bin, bins containing fragments from multiple sources were considered as contaminated, fragments from the same source in different bins were considered as split genomes, and fragments representing an entire source (singletons) were not expected to bin. The following equations are for genome- (source) and bin-based performance metrics, where *B*_*e*_ is the expected number of bins (i.e., sources with at least two fragments), *B*_*g*_ is the number of bins generated, *G*_*e*_ is the expected number of binned fragments (i.e., fragments representing *B*_*e*_ sources), *B*_*o*_ is the total number of bins containing a single source, *G*_*t*_ is the total number of fragments binned, *G*_*b*_ is the number of unique sources binned, *G*_*o*_ is the number of sources contained in a single bin, *G*_*s*_ is the total number of singletons, and *G*_*p*_ the number of binned singletons.

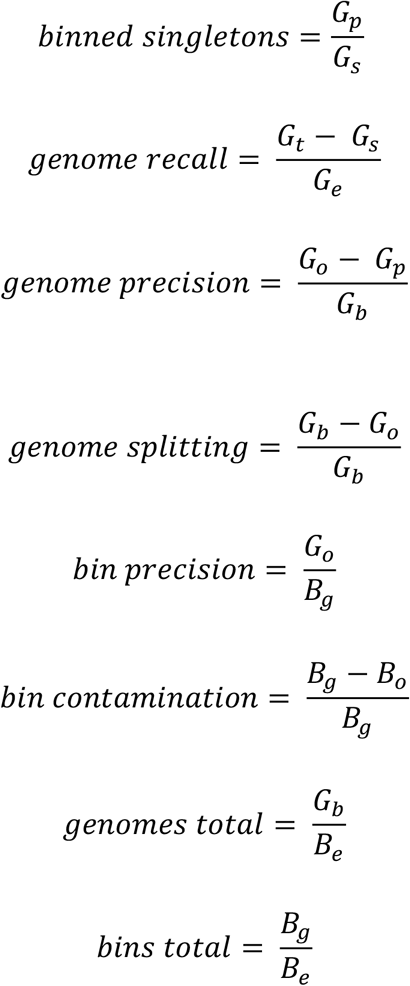

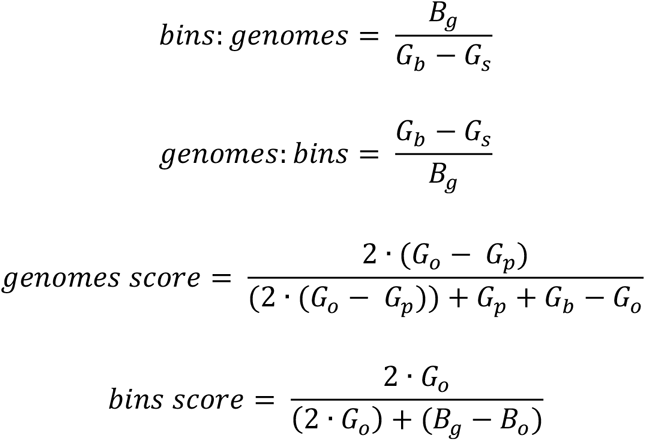

To validate binning further, each pairwise connection between fragments within a bin was evaluated according to each fragment’s nucleotide length. These standard performance metrics were evaluated per bin using true positive *TP*, true negative *TN*, false positive *FP*, and false negative *FN* connections. The following equations are for pairwise nucleotide-based performance metrics:

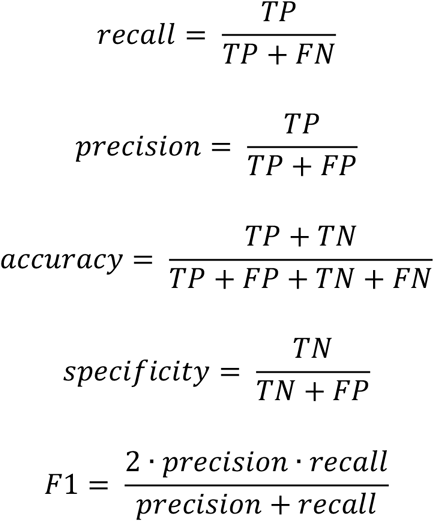

### Performance benchmarking

The performance of vRhyme (v1.0.0) was compared to MetaBat2^15^ (v2.12.1, -s 4000 -m 2000), CONCOCT^26^ (v1.0.0, -l 2000), VAMB^16^ (v3.0.2, -i 2 -m 2000 -t 40), and BinSanity^22^ (v0.5.4, -x 2000). Additional binning tools, namely MaxBin2^14^, MyCC^17^, SolidBin^18^ and DASTool^20^, perform microbial single copy gene analysis and were not applicable, or did not function, for viruses. For VAMB, the starting batch size had to be adjusted to accommodate the relatively small size of the input datasets, and all but three datasets failed to run. The coverage tables for each of the tools were generated from sorted BAM files using each tool’s respective method, except for VAMB for which the same coverage table as MetaBat2 was used. The sorted BAM files were generated using Samtools (v1.13) with reads quality filtered by Sickle^56^ (v1.33) aligned by Bowtie2 (v2.3.5.1, --no-unal --no-discordant).

### Metagenomic datasets and analyses

Publicly available metagenomes from marine^32^, agricultural soil^33^, and human skin^28^ environments were used. Details on the studies, reads used, and accession numbers can be found in **Supplemental Table 11**. Viruses were predicted from each metagenome using VIBRANT and only the identified virus scaffolds were binned using vRhyme. For the human skin datasets, 270 metagenomes from a cohort of 34 individuals with eight body sites per individual were used (antecubital fossa (Af), alar crease (Al), back (Ba), nare (Na), occiput (Oc), toe-web space (Tw), umbilicus (Um), and volar forearm (Vf)). Reads were filtered for quality, adapters, and host-contamination as described previously^28^ using fastp^57^ (v0.21.0, --detect_adapter_for_pe) and KneadData (v0.8.0). MegaHit^58^ (v1.2.9) was used to generate individual metagenomic assemblies for each sample, corresponding to the microbiome of a particular body site for a specific participant at a given timepoint. After predicting viruses, all viruses per body site were combined and dereplicated (--method longest) before binning.

It is important to note that for bins, scaffolds had to be linked with Ns in order to run CheckV analysis since there is no mode to input bins. For all benchmarking using CheckV, the tool was modified to run Prodigal with the -m flag to accommodate linking vMAGs and not predicting open reading frames across the appended strings of Ns connecting scaffolds. For taxonomy of the validation dataset, a publicly available custom reference database of NCBI viruses was used as previously described^59^. In brief, DIAMOND^60^ (v0.9.14) BLASTp^61^ (v2.6.0) was used to identify the most likely taxonomic affiliation of a sequence.

### Additional datasets and benchmarking

Additional publicly available datasets were used to assess the performance of vRhyme under different scenarios and conditions. To assess binning of related types of viruses within the same sample, a total of 101 publicly available crAssphage sequences^62^ were dereplicated using vRhyme (--derep_id 0.97 --frac 0.70 --method longest) to 86 non-redundant scaffolds. The non-redundant scaffolds were randomly fragmented as described previously into 791 fragments. To assess binning of megaphages and eukaryotic viruses with large genomes, the 540 kb Prevotella phage Lak C1^63^ was randomly fragmented into 51 fragments, and four different eukaryotic viruses^64,65^ with genome lengths ranging from 154 kb to 201 kb were each randomly fragmented into 11 to 19 fragments. To assess binning of active and dormant prophages, VIBRANT was used to predict prophage regions for 10 active prophages from 3 different hosts and 24 dormant prophages from 5 different hosts. Activity or dormancy was determined according to respective studies described elsewhere^66–68^ and validated using PropagAtE^69^ (v0.9.0). Whole prophage scaffolds from the same host genome were binned together. Details on the studies, reads used, scaffolds, and accession numbers can be found in **Supplemental Table 11**.

To validate protein redundancy, NCBI databases (RefSeq and Genbank, release July 2019) were queried for “prokaryotic virus” as before and genomes greater than 3 kb in length were retained. Likewise, NCBI databases (RefSeq and Genbank, release September 2021) were queried for “eukaryotic virus” and genomes greater than 20 kb in length were retained. Proteins were predicted using Prodigal (-p meta) for 15,238 prokaryotic and 557 eukaryotic viruses. Protein redundancy was calculated per genome based on the method described for vRhyme, with the exception that proteins could be redundant if encoded along the same scaffold.

### Effect of number of samples

The effect of the number of input samples on vRhyme performance was done by stepwise increasing the number of BAM files used to calculate coverage from one to the maximum number of samples for a given dataset. To do this, samples were arranged in descending order, starting at the sample with the greatest total coverage across all scaffolds and were stepwise combined, ending with the sample with the lowest coverage.

### Visualizations

All plots and visualizations were done using Matplotlib^70^ (v3.2.0) and Seaborn^71^ (v0.11.0). Genome alignment visualizations were made using EasyFig^72^ (v2.2.2) and Geneious Prime 2019.0.3. Genome alignments to identify percent sequence identity were made using progressiveMauve^73^ (development snapshot 2015-02-25). vConTACT2^30^ (v0.9.19, --rel-mode Diamond --db ‘None’ --pcs-mode MCL --vcs-mode ClusterONE, ClusterONE^74^ v1.0) was used to construct protein clustering networks and visualized using Cytoscape^75^ (v3.7.2).

## Code availability

vRhyme and all auxiliary scripts are freely available as open-source Python code at https://github.com/AnantharamanLab/vRhyme.

## Acknowledgements

We thank members of the Anantharaman laboratory at the University of Wisconsin-Madison for helpful feedback and discussions. This research was supported by National Institute of General Medical Sciences of the National Institutes of Health under award number R35GM143024 to K.A. and award numbers R35GM137828 and U19AI142720 to L.K. A.A. was funded by a University of Wisconsin-Madison CIBM postdoctoral traineeship from the National Library of Medicine (T15LM007359). K.K. was supported by a Wisconsin Distinguished Graduate Fellowship Award from the University of Wisconsin-Madison, and a William H. Peterson Fellowship Award from the Department of Bacteriology, University of Wisconsin-Madison.

## Author contributions

K.K. and K.A. designed the study. K.K., A.A., and R.S. developed code and conducted bioinformatic analyses. K.K. and K.A drafted the manuscript. All authors (K.K., A.A., R.S., L.K., and K.A.) reviewed the results and approved the manuscript.

