## Supplementary Text for "vRhyme enables binning of viral genomes from metagenomes"

*Benchmarking vRhyme on NCLDV, megaphage, large eukaryotic viruses, crAssphage, integrated prophages and microbes*

We next evaluated the capability of vRhyme to handle large genomes, and high similarity or complex datasets. First, to validate that vRhyme's metrics are applicable to large viral genomes, we binned a 540 kb megaphage that was split into 51 randomly sized fragments. Using a single coverage sample[1], vRhyme was capable of binning all 51 fragments into a single bin. Additionally, we utilized a publicly available single-sample metagenome containing a giant ~1.6 Mb virus, specifically a Fadolivirus-like giant nucleocytoplasmic large DNA virus (NCLDV). Originally, the NCLDV was identified as a 12-scaffold bin generated by MetaBat2[2], in which only 11 of the scaffolds aligned to a Fadolivirus reference genome[3]. Using the entire metagenome containing 89,569 scaffolds as input, vRhyme generated a respective NCLDV bin, representing only the 11 scaffolds that align to the reference genome. As an additional validation, we evaluated protein redundancy for both the megaphage and the NCLDV. As expected, the megaphage contained no redundant proteins. The NCLDV bin contained 9 redundant proteins, though these are more anticipated as larger eukaryotic viruses (e.g., Pandoraviridae, Mimiviridae, Herpesviridae, Pithoviridae) tend to encode redundant proteins based on vRhyme metrics.

Next, vRhyme was tested on four dsDNA eukaryotic viruses: African swine fever virus, Camelpox virus, Human gammaherpesvirus 8, and Molluscum contagisum virus (**Supplementary Table 5a**)[4,5]. The viral genome sizes range from 154 kb to 201 kb and were each randomly fragmented into 11 to 19 fragments. Binning with vRhyme, with one sample read set per virus, yielded recalls of 82% to 95% based on the number of fragments binned per genome. For three of the viruses a single bin was generated, indicating high precision, whereas one of the viruses was split into 3 separate bins. We also combined the fragments from all of the viruses together and binned using all four read sets as the input for coverage. vRhyme yielded identical results as when the viruses were binned individually. This is as expected, however, since differential coverage per virus across the four samples was distinct. We did not test vRhyme on small DNA or RNA eukaryotic viruses as these are less likely to require binning.

Furthermore, we evaluated if vRhyme could handle an input of many viral genomes with similar evolutionary similarities, namely a shared host range and phylogeny. To do this, 86 non-redundant crAssphage metagenome-derived genomes[6] with varying completeness were randomly fragmented into 791 fragments (**Supplementary Tables 5b,c**). Recall was relatively low (0.45) in this high similarity dataset, though precision (0.94) and specificity (0.97) remained high. A genome score of 0.65 and a bin score of 0.98 were calculated, which indicates that although several genomes were split across multiple bins, most bins were accurate. A total of 95 bins were generated representing 60 unique genomes with 55 redundant proteins identified across all bins, but 53 of these proteins originated from only two bins that each were a mix of fragments from the same two genomes. In fact, only 3 of the 95 bins were contaminated, and of those 2 were readily identified as contaminated based on protein redundancy. Furthermore, the two highly contaminated bins were comprised of fragments from genomes with >99.9% nucleotide identity across 54 kb and likely represent the same crAssphage population. Based on this, we conclude that vRhyme is highly accurate in handling datasets comprised on similar viral populations. Although many viral genomes in this example were split across multiple bins, vRhyme still represents a significant improvement over foregoing binning, especially since the resulting bins are precise. This is further supported by binning the complete scaffolds of the 86 crAssphages together with the expectation that none should bin. Here, only a single bin was generated, comprised of the exact same two scaffolds with >99.9% nucleotide identity described above. In comparison, MetaBat2 binned 57 scaffolds into 16 bins.

Lastly, we evaluated binning of prophages integrated within the same host genome. To do this we evaluated active and dormant prophages. Details of which viruses were active or dormant per host as well as the binning results can be found in **Supplementary Table 5d**. Here, differential coverage is essential for binning due to high similarities in nucleotide markers in prophages integrated within the same host. Active prophages will likely have genome coverage distinct from that of their host as well as other prophages and dormant prophages will have uniform coverage with both the host genome and other prophages. We evaluated 24 dormant prophages[7–11] within *Clostridium limosum*, *Kosakonia* sp., *Lactococcus lactis*, *Metakosakonia* sp. and *Sphingomonas paucimobilis*, and 10 prophages integrated within *Bacillus licheniformis*, *Bartonella krasnovii* and *Lactococcus lactis* genomes, of which 4 were active[12–14]. The 2 dormant prophages of *Clostridium limosum*, 3 of *Sphingomonas paucimobilis* and 10 of *Metakosakonia* sp. each generated one bin. Similarly, the 6 dormant prophages of *Lactococcus lactis* generated two bins. Only the 3 dormant prophages of *Kosakonia* sp. did not generate a bin. This indicates a significant caveat in the binning of viruses, in that dormant integrated prophages with similar coverage profiles within the same host typically yield a single, combined bin. However, a distinction of active prophages could be made. The 1 active prophage of *Bartonella krasnovii* and 1 of *Lactococcus lactis* each did not bin with the other 3 dormant prophages in the respective host genome. Likewise, the 2 active prophages of *Bacillus licheniformis* did not bin together. Overall, it is likely that active prophages will generate uncontaminated bins, whereas dormant prophages with indistinct coverage profiles will bin poorly.

In additional to viral datasets, we also checked if vRhyme could bin non-viral genomes. We anticipated that vRhyme would perform poorly on datasets comprised mainly of non-viral scaffolds since it was not developed for this purpose. For this we chose to bin a CAMI medium complexity dataset[15], limited to only sequences at least 2 kb, and compare the results to MetaBat2. As anticipated, vRhyme binning resulted in a lower F1 score (0.74) compared to MetaBat2 (0.83), mainly due to lower specificity (0.45 versus 0.61, respectively) (**Supplementary Table 6**).

*Benchmarking vRhyme on marine viromes*

We next applied vRhyme to the Global Ocean Virome 2 (GOV2) database[16] and compared the results to MetaBat2. For metagenomic datasets such as GOV2 the expected number of scaffolds to bin and the number of bins is unknown. First, all scaffolds from the GOV2 database were limited to scaffolds at least 5 kb in length and dereplicated by 98% identity. Of the 108,947 input scaffolds, vRhyme binned 56,642 scaffolds into 13,175 bins and MetaBat2 binned 57,800 scaffolds into 11,826 bins. Despite the number of scaffolds binned being relatively similar, the number of bins generated was quite different. However, vRhyme yielded 15,106 redundant proteins whereas MetaBat2 yielded 29,334, indicating that vRhyme was likely more precise and generated fewer contaminated bins (**Supplementary Figure 3a**). In support of this, vRhyme generated 1,266 bins with at least 2 redundant proteins and MetaBat2 generated 1,648. When these likely contaminated bins were removed, vRhyme binned 48,251 scaffolds into 11,909 bins and MetaBat2 binned 33,351 scaffolds into 10,178 bins (**Supplementary Figure 3b**). Based on protein redundancy, vRhyme was capable of binning far more viral scaffolds and generating more bins of low contamination compared to MetaBat2. Note, we identified bins with "low contamination" to be 0-1 redundant proteins based on a benchmark of prokaryotic and eukaryotic viral genomes from NCBI databases (**Supplementary Figure 4**).

We also estimated the completeness of the 11,909 low contamination vRhyme bins using CheckV (**Supplementary Figures 3c,d**). Contamination was not estimated using CheckV as that metric does not consider contamination of multiple viral genomes, but rather contamination of non-viral sequences. The binned scaffolds individually yielded 25,969 completeness values with an average of 14% complete, 79 estimated to be 100% complete, and 22,282 with 'NA' completeness. The scaffolds within each bin were then linked into vMAGs, which yielded 8,393 completeness values with an average of 48% complete, 775 estimated to be 100% complete, and 3,516 with 'NA' completeness. Therefore, vRhyme generated vMAGs with greater average completeness to aid in downstream analyses and interpretations, even with high complexity or larger datasets such as GOV2.

*Benchmarking vRhyme on agricultural soil viromes*

Next, we applied vRhyme, MetaBat2 and VAMB to agricultural soils, which were shown to harbor highly diverse and complex viral communities. A total of 3,690 viral scaffold from two virome studies (15 and 7 samples) and one metagenome study (16 samples) were used[17]. Without filtering for contamination by protein redundancy all three tools performed relatively poorly (**Supplementary Tables 7a**). In the two virome sets of samples, MetaBat2 binned more scaffolds (2,381 and 2,424) compared to vRhyme (1,853 and 2,284) and VAMB (2,200 and 1,952) but at a cost to contamination in the form of redundant proteins (13,345 and 12,868) compared to vRhyme (12,496 and 9,292) and VAMB (13,518 and 12,854). MetaBat2 also generated more bins (875 and 801) than vRhyme (699 and 722) and VAMB (804 and 727) but for both MetaBat2 and VAMB those bins on average were more contaminated. The results were similar for the metagenome study, in which MetaBat2 and vRhyme binned relatively equal number of scaffolds (224 and 254, respectively), fewer than VAMB (837), but vRhyme generated fewer bins than MetaBat2 (66 and 76, respectively) with lower contamination (685 and 868 redundant proteins, respectively). VAMB generated more bins (333) than MetaBat2 and vRhyme along with much higher contamination (6,427).

Due to the high degree of contamination, we next filtered by low contamination (0-1 redundant proteins per bin) bins for a more competitive comparison (**Supplementary Table 7b**). In the first virome set, the number of scaffolds binned and number of bins generated by vRhyme (673 and 291, respectively), MetaBat2 (1003 and 421, respectively) and VAMB (929 and 382, respectively) were dissimilar, with MetaBat2 appearing to perform optimally. Yet, it is possible that vRhyme, despite binning fewer scaffolds, generated fewer bins of higher quality. In the second virome set, vRhyme (1151 and 412) and MetaBat2 (1015 and 391) binned relatively equal number of scaffolds and total bins, respectively, with VAMB binning slightly less (751 and 322). The same was true for the metagenome study, though VAMB binned slightly more scaffolds. In all cases, there was no difference in total redundant proteins after filtering by contamination.

Based on the results of binning agricultural soils, vRhyme was capable of handling highly complex and diverse input viral scaffolds. However, the bins required refinement, based on information provided by vRhyme from binning, as the raw results contained many bins with redundant proteins as a signature of contamination. The performance of vRhyme after filtering the bins by contamination was comparable to MetaBat2 and VAMB with no tool distinctly performing more optimally across all the datasets.

*Effect of coverage algorithm*

We compared the performance of vRhyme on three of the nine performance benchmarking datasets using the coverage table generated from the native coverage calculation method to the coverage table generated by MetaBat2 (**Supplementary Table 8**). For this, vRhyme comes with an auxiliary Python script that can convert a MetaBat2-generated coverage table into vRhyme format. In all three datasets, the performance of vRhyme marginally increased F1 scores by 0.01 to 0.04 with the MetaBat2 coverage table. However, the genome scores and bin scores remained largely unchanged with a maximum increase of 0.01 with the MetaBat2-generated coverage table for genome score in one dataset. Therefore, the native coverage calculation implemented by vRhyme appears to be robust in comparison to the state-of-the-art method implemented by MetaBat2.

*Effect of samples*

The performance of vRhyme on a given set of input sequences is dependent on two factors: the number of samples for coverage analysis and the set parameters. A common rule is that binning performance increases with the addition of samples due to higher statistical confidence in calculating co-occurring scaffolds, with an optimal minimum of three samples.

We evaluated the effect of the number of input samples on binning performance using three datasets: GOV2[16] (55 samples), crAssphages (171 samples), and a freshwater dataset[18] (12 samples) (**Supplementary Table 9, Supplementary Figure 5**). To do this, for each dataset, the samples were arranged in descending order by total coverage across the scaffolds. Binning started with one input sample, representing the sample with the greatest total coverage, and samples were stepwise added. The crAssphage and freshwater dataset were evaluated based on expected performance as described above whereas GOV2 was evaluated based on the raw binning results. For GOV2 and the freshwater dataset, binning performance increased significantly from one to three input samples, but for the crAssphges performance did not increase noticeably until six samples. For all three datasets, performance leveled out after approximately 40-60% of the samples were added. For GOV2 and the freshwater dataset, the performance change after this mark was insignificant, though surprisingly led to a minor decrease in precision. This observation was starker with the crAssphages, in which F1 score decreased from 0.67 at 66 samples to 0.59 at 105 samples and remained unchanged until 171 samples. The performance decrease was likely due to the addition of confounding samples, in which low and uneven coverage across scaffolds leads to statistical noise. Taken together, a single input sample is sufficient to yield results, though a general minimum of at least 3 samples will be more likely to yield results of higher quality. However, this threshold is dependent on the manner in which the dataset was created. For example, datasets generated from greater than 10 samples that contain many scaffolds with low coverage may have optimal performance when only ~40-60% of the samples are utilized, starting with those with the greatest total coverage.

*Effect of parameters*

The parameters that most effect vRhyme are coded with the iteration presets and are immutable. Therefore, the default settings for vRhyme are likely sufficient due to the iteration over multiple, optimal preset parameters. We evaluated the effect of vRhyme parameters on binning performance, based on F1 score, for five datasets. It is important to note, however, that the resulting effect of parameter changes in this analysis will differ compared to metagenomes. Since vRhyme allows the selection of alternative bins from any iteration, we assessed the ten default iteration presets individually but found that despite only minor changes in F1 score, the most optimum preset was selected as the best iteration. For the five datasets tested, setting the number of iterations (iter) from 8 to 15 had no effect on performance due to the most optimized presets being set within the first 8 iterations (**Supplementary Figure 6a,b**).

We next evaluated parameters effecting coverage calculations: minimum coverage (cov), scaffold end masking (mask), coverage penalty weight (penalty_w), coverage penalty number (penalty_n), and minimum read identity (read_id) (**Supplementary Figures 6c-g**). For each, the default settings generated the best performance. Minimum coverage, penalty weight and read identity had identifiable effects on performance, but only if modified relatively far from the default settings. Scaffold end masking had no effect on these datasets and penalty number only had a significant effect if set to zero. The effect of combining parameter changes, such as increasing read identity while decreasing minimum coverage, was not evaluated due to the large number of possible combinations.

The remaining parameters effect nucleotide comparisons and bin generation: minimum kmer distance (min_kmer), maximum GC distance (max_gc), the machine learning model(s) used (model), maximum network edges (max_edges), minimum bin size for refinement (mems), and maximum redundancy per bin (red) (**Supplementary Figures 6h-m**). Increasing the minimum kmer distance beyond the default results in a significant decrease in performance by F1 score due to low recall, though precision is likely to increase. Using the hybrid approach for machine learning, in which two models are used in tandem, usually leads to the best performance but only marginally. For the remaining parameters, the default settings only need to be modified for specific functionality, such as setting maximum redundancy to 0%, though these filters can be applied post-binning.

*Runtime, memory and reproducibility*

vRhyme was optimized for high-speed performance and a low memory burden in order to efficiently process metagenome samples. The runtime and max memory usage of vRhyme was benchmarked on several datasets of varying sizes. The first benchmarks are for implementation of the binning functionality and do not include read mapping or generation of the input coverage table, though the latter two can be automated within the vRhyme pipeline. For average size datasets with approximately 500 to 4000 sequences vRhyme completed binning in under one minute (8 – 40 seconds) using five CPU threads. Even with a dataset of 3,690 sequences, 7 coverage samples, and 1 CPU thread vRhyme completed binning in 86 seconds. The max memory usage for all of these benchmarks was ~400 MB, which is primarily due to loading the machine learning model. We also benchmarked the runtime of generating the coverage table from sorted BAM files. For the same three datasets, the coverage tables also generated quickly (0.5 – 3.6 minutes), with the longest runtime of 3.6 minutes for a dataset with 2,130 scaffolds, 96 BAM files and five CPU threads. The CAMI medium dataset was binned with 10 CPU threads and 0.71 GB memory in 4 minutes (**Supplementary Table 10**).

Furthermore, for the GOV2 dataset with 108,947 sequences and 55 coverage samples, vRhyme completed binning in 93 minutes with 15 CPU threads and used 2.3 GB of memory. We also assess the effect of the number of input samples on binning runtime for the full GOV2 dataset. Runtime increased logistically, with a steep runtime increase from 1-17 samples and mostly stable runtimes (88-93 minutes) for 33-55 samples (**Supplementary Figure 7**). Due to how vRhyme processes coverage, the leveling is likely the result of minimal additional coverage variance information provided after 33 samples. Across all samples memory usage remained constant. Overall, we demonstrate that vRhyme has practical computational demands for viral datasets.

As a final assessment, we validated that vRhyme is able to generate reproducible results. In fact, vRhyme reproduces identical results on a given dataset if the inputs remain constant. To show this, we binned each performance dataset multiple times, including the larger GOV2 dataset five separate times. In all instances vRhyme generated identical results.
