## Supplementary Figures 1-7 for "vRhyme enables binning of viral genomes from metagenomes"

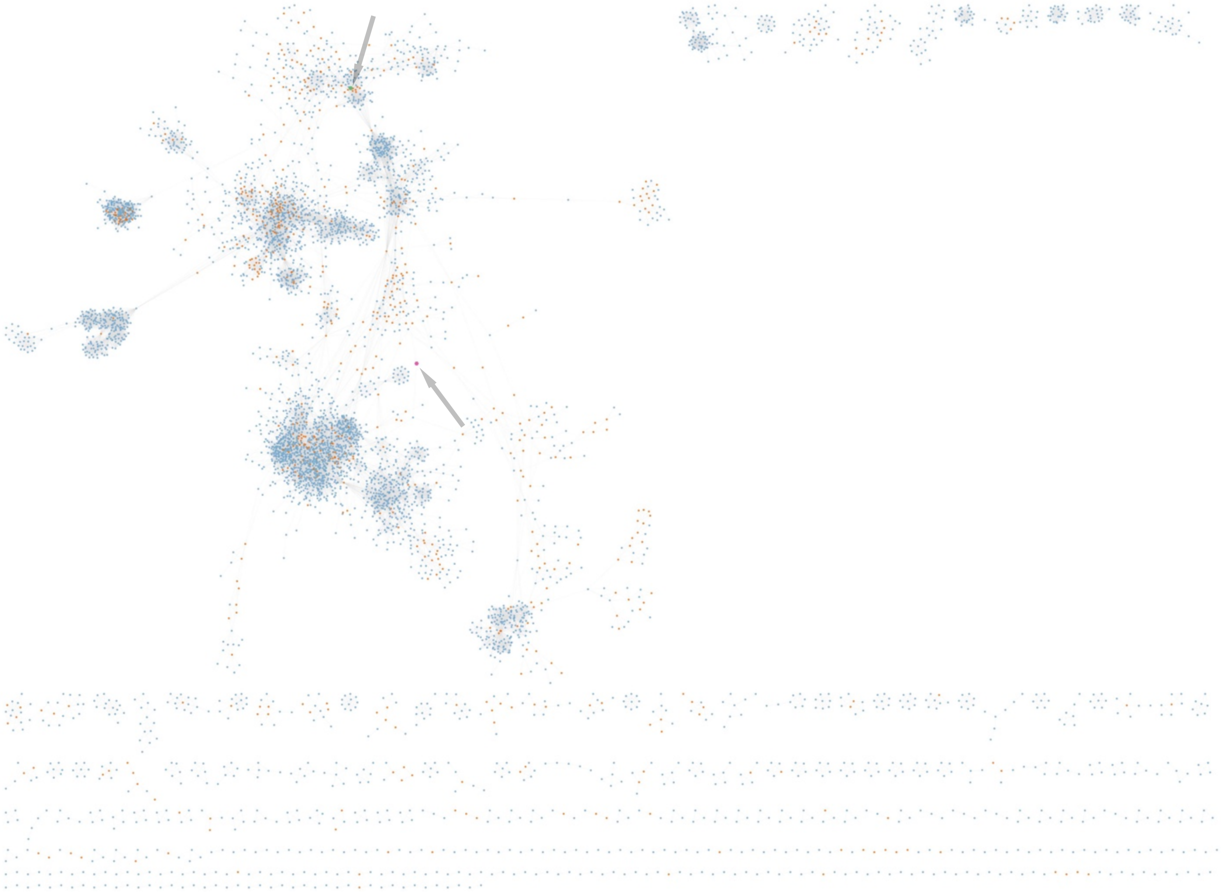

**Supplementary Figure 1.** vConTACT2 network clustering of the complete human skin binning results (vMAGs plus unbinned vOTUs). Blue: vOTUs, orange: vMAGs, green: Vf bin 113 (see arrow), purple: Tw bin 8 (see arrow).

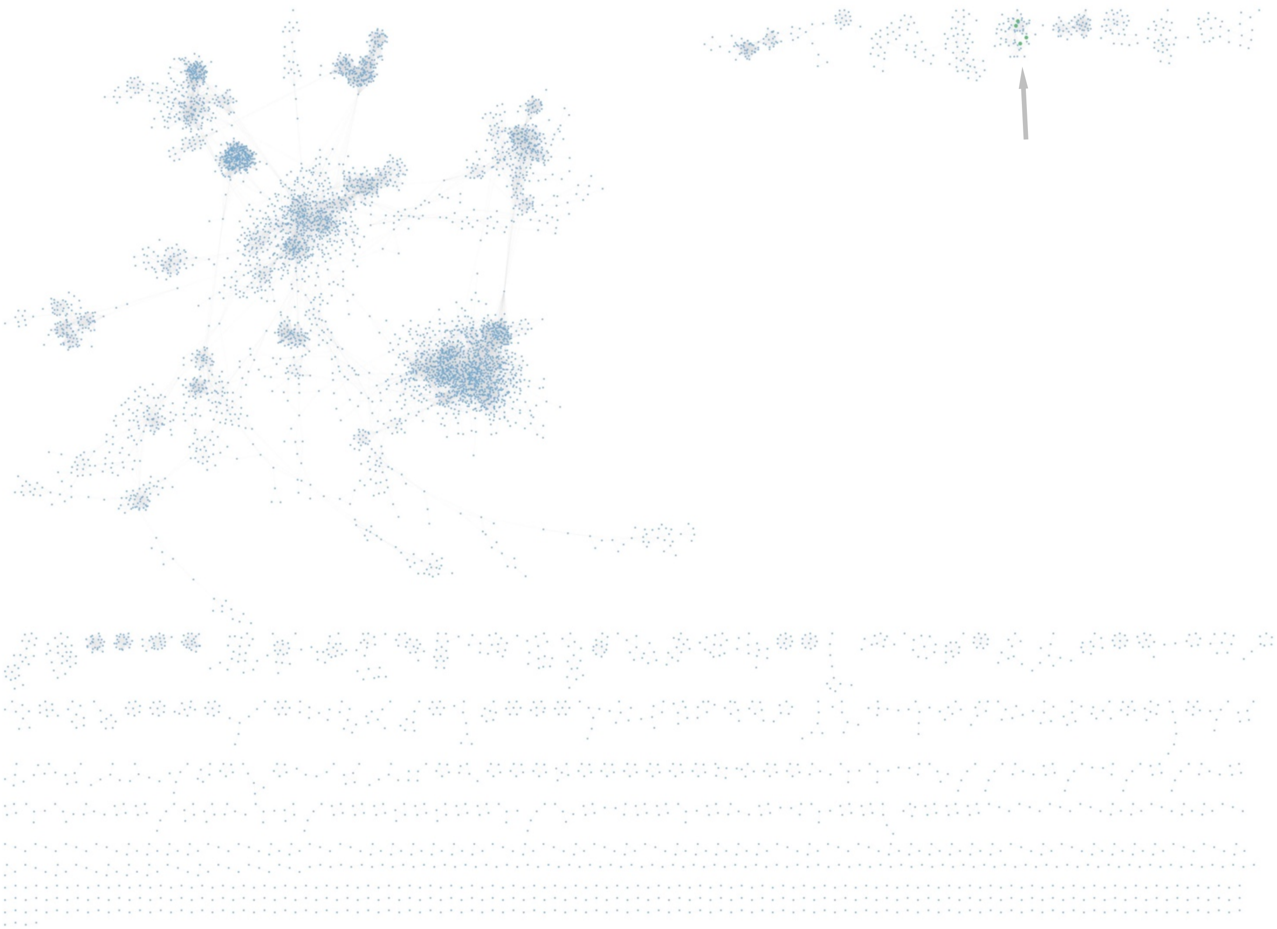

**Supplementary Figure 2.** vConTACT2 network clustering of the human skin individual, unprocessed viral scaffolds. Blue: vOTUS, green: individual scaffolds in Vf bin 113 (see arrow).

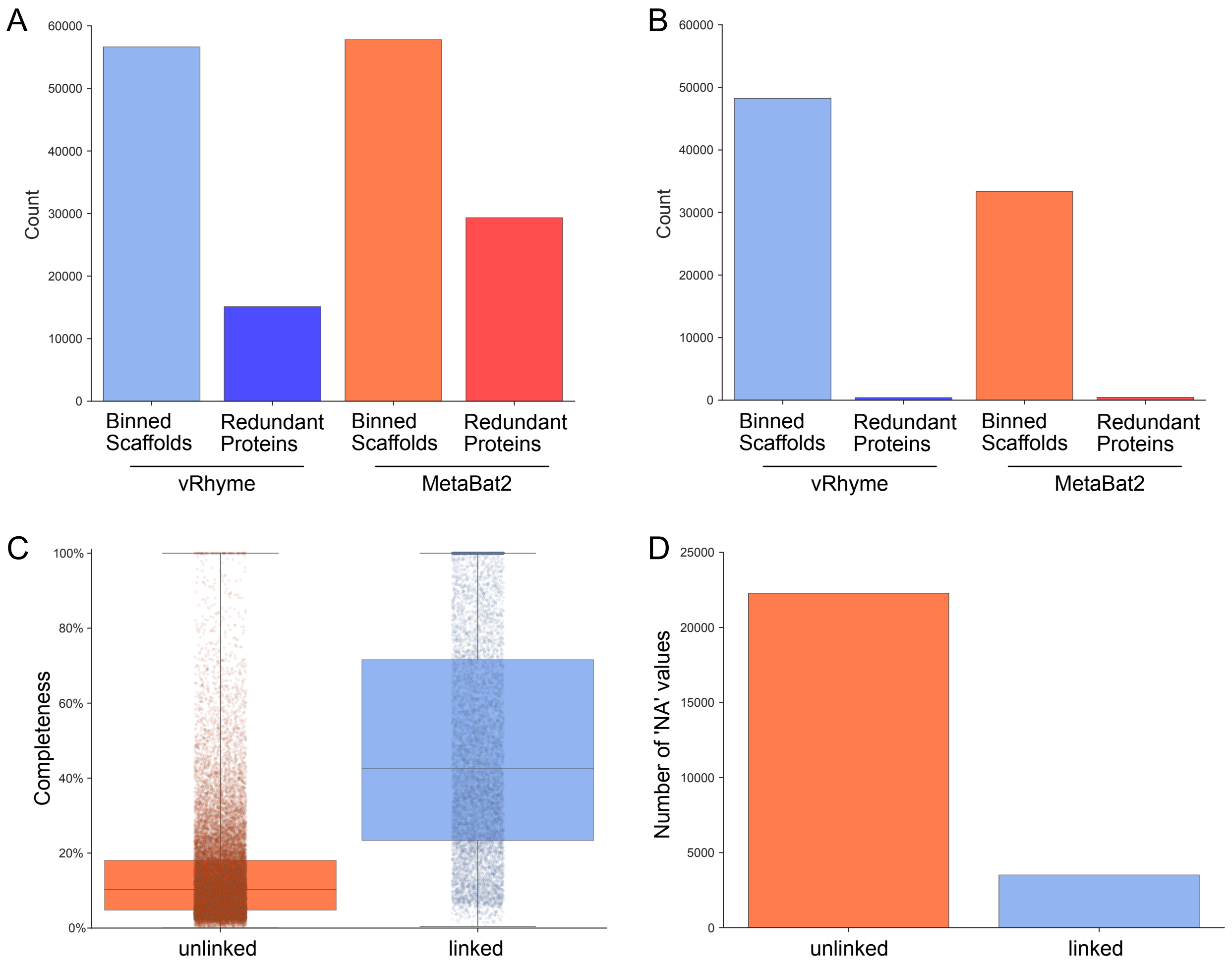


**Supplementary Figure 3. Benchmark binning and completeness evaluation of GOV2**. Comparison of vRhyme and MetaBat2 (**a**) raw results and (**b**) low contamination filtering results by the number of scaffolds binned and identified redundancy. For vRhyme, CheckV was used to identify (**c**) the estimated completeness values and (**d**) number of 'NA' completeness values of the raw results for individual binned scaffolds as well as whole vMAGs.

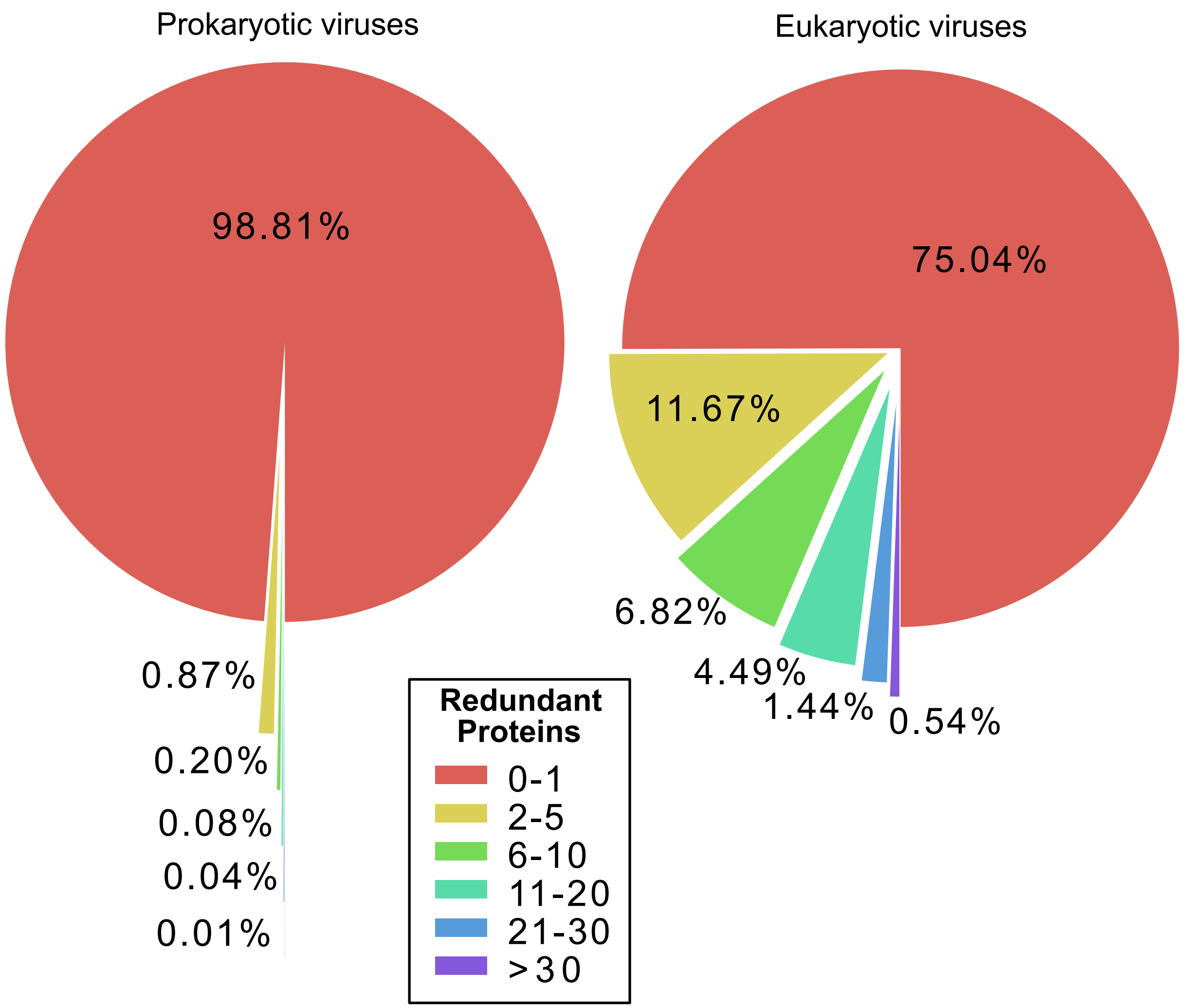


**Supplementary Figure 4. Evaluation of protein redundancy of NCBI reference viruses**. The vRhyme protein redundancy method was used to benchmark complete (**a**) prokaryotic and (**b**) eukaryotic viruses, with the exception that redundant proteins along the same scaffold were considered.

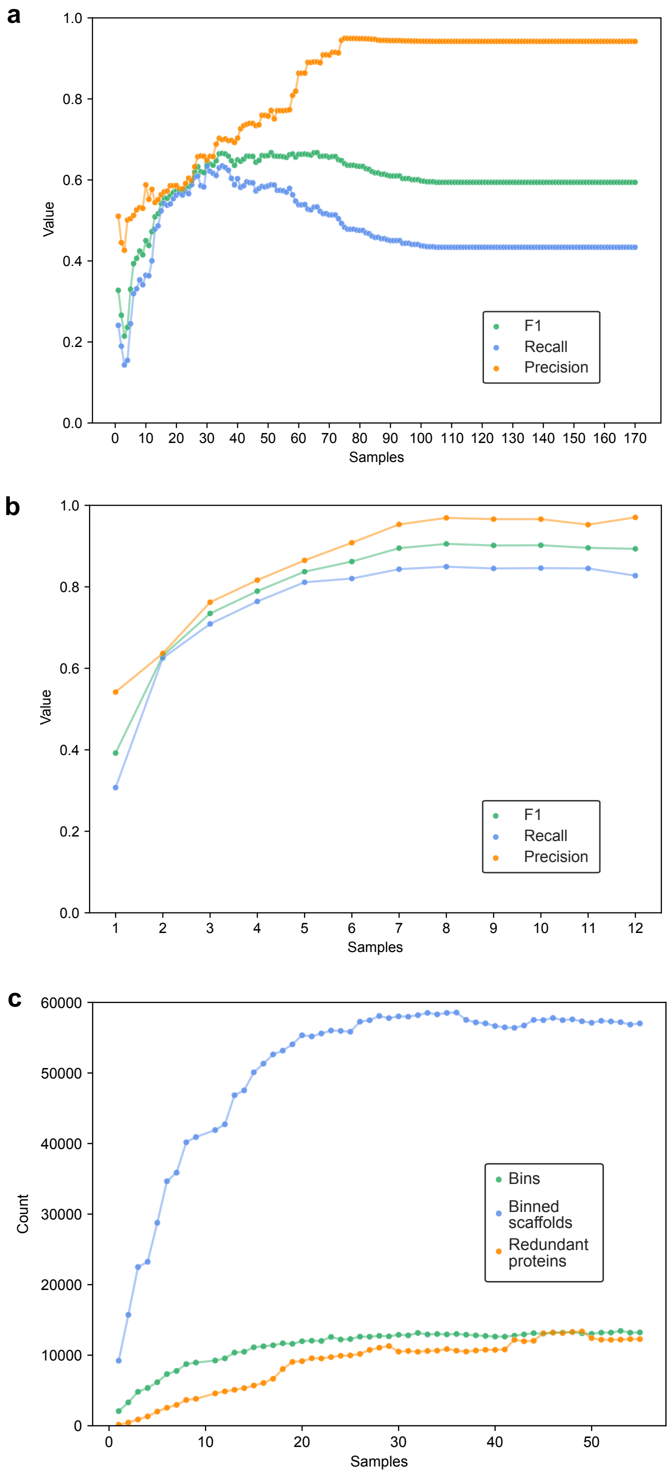


**Supplementary Figure 5. Effect of the number of input samples for read coverage comparisons on vRhyme performance**. F1 score, recall and precision are compared for (**a**) the crAssphage dataset and (**b**) a freshwater dataset. Number of bins, members and redundant proteins are compared for (**c**) the GOV2 dataset. Each dot represents a single sample and the lines connecting samples are for visualization of trends.

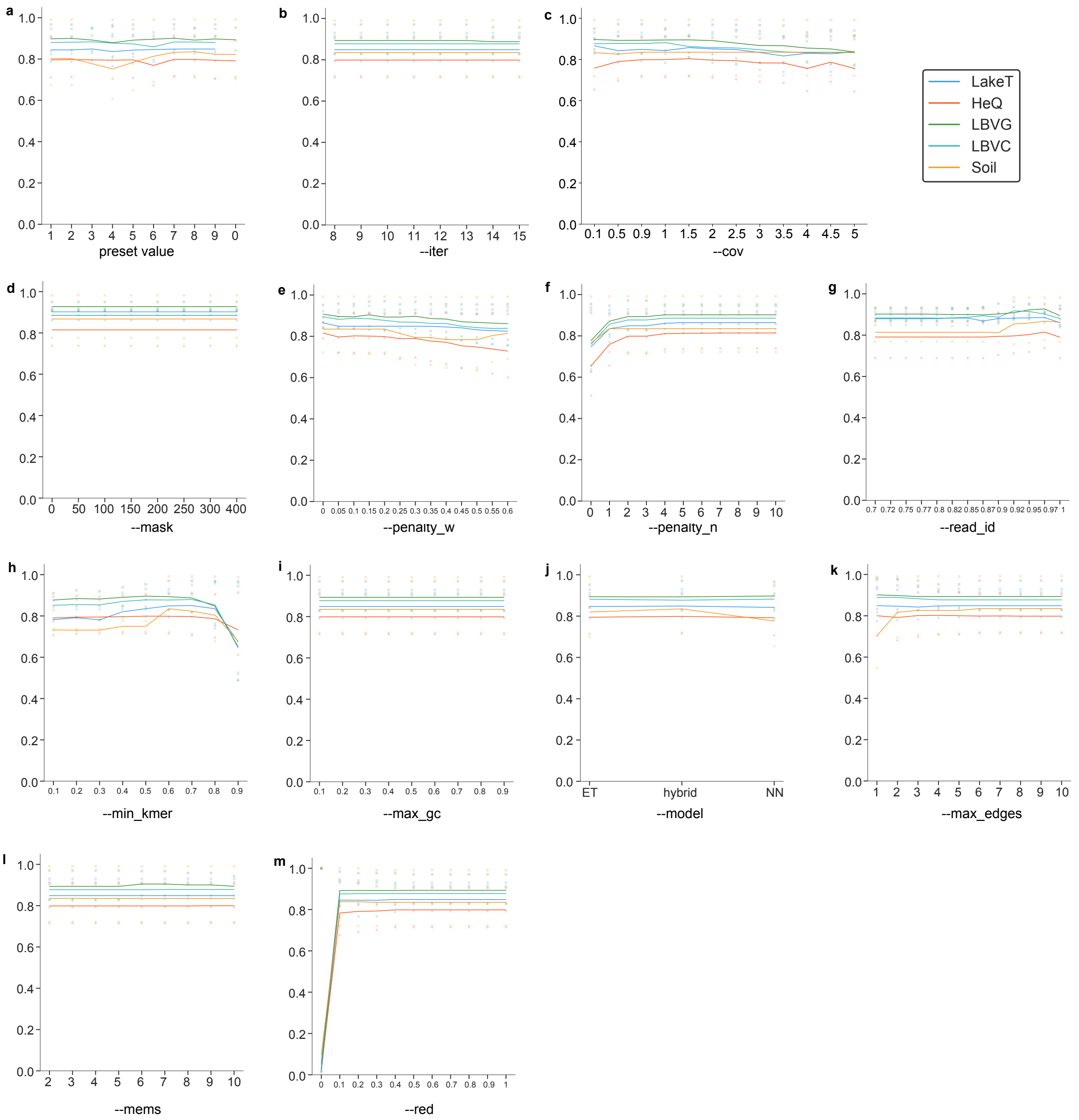


**Supplementary Figure 6. Effect of presets and parameters on vRhyme performance**. For all, F1 score is used to evaluate performance on five separate datasets. **a**, effect of the preset value, which is an unmodifiable parameter. **b-m**, effect of modifiable flags that may impact binning: iter, cov, mask, penalty_w, penalty_n, read_id, min_kmer, max_gc, model, max_edges, mems and red, respectively. Each dot represents a single sample and the lines connecting discrete points are for visualization of trends.

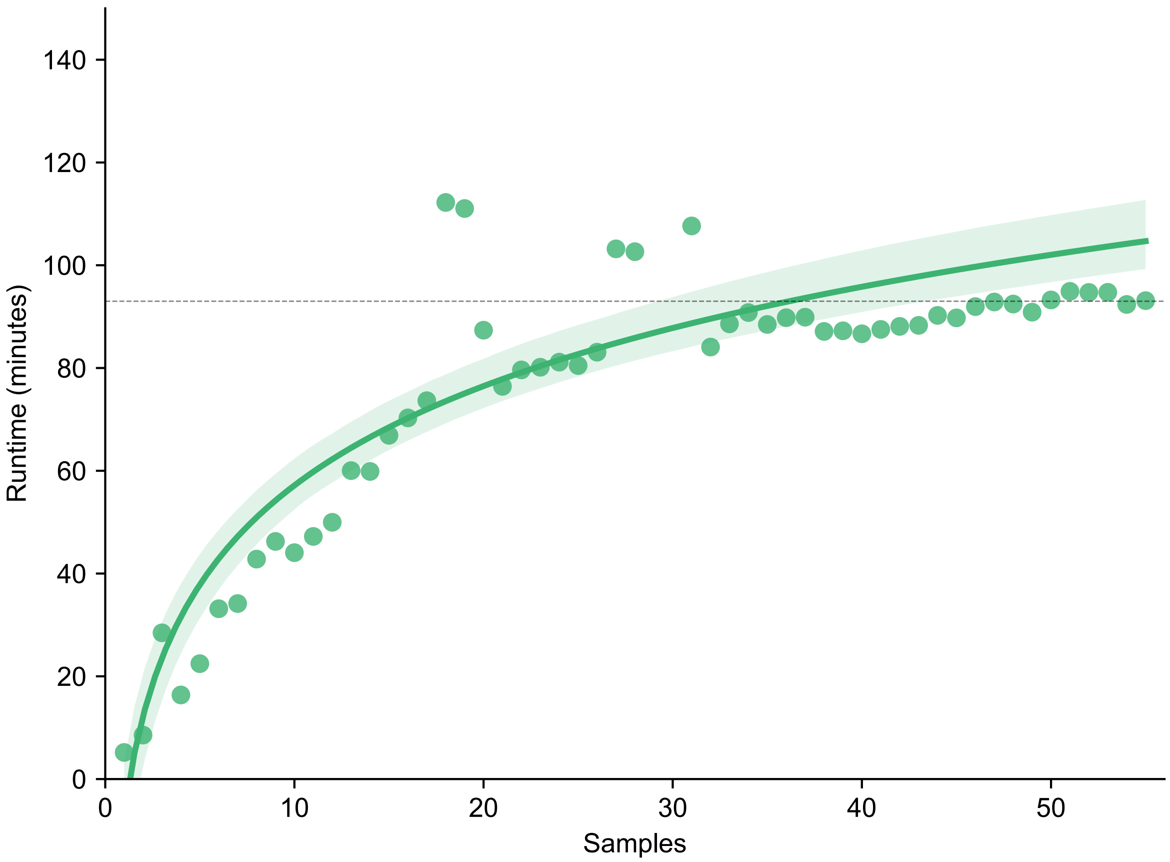


**Supplementary Figure 7. Effect of the number of samples on vRhyme runtime.** GOV2 was binned by stepwise adding samples for read coverage comparison from 1-55. The runtime in minutes was evaluated using 15 CPU threads. A logarithmic trendline was fitted to the data was a 95% confidence interval (shading) and a dotted line at 93 minutes shows the runtime for 55 samples. Variations in runtime near the center likely represent fluctuations in computer performance. For each, the maximum memory usage was 2.23 GB.
